# Heritable Genetic Background Alters Survival and Phenotype of *Mll-AF9*-Induced Leukemias

**DOI:** 10.1101/2020.06.13.149310

**Authors:** Kira Young, Matthew A. Loberg, Elizabeth Eudy, Logan S. Schwartz, Kristina D. Mujica, Jennifer J. Trowbridge

**Affiliations:** The Jackson Laboratory for Mammalian Genetics, Bar Harbor, ME 04609

## Abstract

The MLL-AF9 fusion protein occurring as a result of t(9;11) translocation gives rise to pediatric and adult acute leukemias of distinct lineages, including acute lymphoblastic leukemia (ALL), acute myeloid leukemia (AML), and mixed phenotype acute leukemia (MPAL). The mechanisms underlying how this same fusion protein results in diverse leukemia phenotypes among different individuals is not well understood. Given emerging evidence from genome-wide association studies (GWAS) that genetic risk factors contribute to *MLL*-rearranged leukemogenesis, here we tested the impact of genetic background on survival and phenotype of a well-characterized *Mll*-*AF9* knockin mouse model. We crossed this model to five distinct inbred strains (129, A/J, C57BL/6, NOD, CAST), and tested their F1 hybrid progeny for dominant genetic effects on *Mll-AF9* phenotypes. We discovered that genetic background altered peripheral blood composition, with *Mll-AF9* CAST F1 demonstrating significantly increased B lymphocyte frequency while the remainder of the strains exhibited myeloid-biased hematopoiesis, similar to the parental line. Genetic background also impacted overall survival, with *Mll-AF9* A/J F1 and *Mll-AF9* 129 F1 having significantly shorter survival, and *Mll-AF9* CAST F1 having longer survival, compared to the parental line. Furthermore, we observed a range of hematologic malignancies, with *Mll-AF9* A/J F1, *Mll-AF9* 129 F1 and *Mll-AF9* B6 F1 developing exclusively myeloid cell malignancies (myeloproliferative disorder (MPD) and AML) whereas a subset of *Mll-AF9* NOD F1 developed MPAL and *Mll-AF9* CAST F1 developed ALL. This study provides a novel *in vivo* experimental model to evaluate the underlying mechanisms by which *MLL-AF9* results in diverse leukemia phenotypes and provides definitive experimental evidence that genetic risk factors contribute to survival and phenotype of *MLL*-rearranged leukemogenesis.

## Introduction

Chromosomal rearrangements involving the *mixed-lineage leukemia 1* (*MLL1*) gene, also known as *Lysine [K]-specific methyltransferase 2A* (*KMT2A*), generate fusion proteins causing aggressive acute leukemias in infants, children and adults. *MLL*-rearranged leukemias comprise ∼10% of acute leukemias across all age groups^1^. Patients with *MLL-*rearranged leukemias generally have a poor prognosis, with high-risk treatment options and frequent relapse. This underscores an unmet need for novel therapeutic approaches to improve outcomes in *MLL-*rearranged leukemia.

Emerging evidence from GWAS studies suggest that heritable genetic polymorphisms can modify the risk of *MLL*-rearranged leukemia^2-4^. To definitively test causation and build upon these findings toward development of novel therapeutic targets, use of *in vivo* mouse models of *MLL*-rearranged leukemia is ideal. However, the vast majority of genetically engineered mouse models of human leukemia are studied on a single inbred genetic background, C57BL/6, despite genetic variability having been recognized as an important modifier of leukemogenesis in mouse models^5-8^.

The first MLL fusion protein to be modeled as an endogenous knockin allele in mice was *Mll-AF9* (t(9;11))^9^. After an early myeloproliferative phase, *Mll-AF9* mice primarily succumb to AML, and only in rare cases to ALL^10,11^. This is notably distinct from human disease, where *MLL-AF9* is found in both B-cell ALL (B-ALL) and AML in infants and children, and AML in adults^12^. Here, we have utilized this well-characterized *Mll-AF9* knockin mouse model to test the extent to which dominant genetic alleles modify *Mll-AF9-*driven leukemogenesis using genetically diverse mouse strains^13,14^.

## Results and Discussion

To determine the role of genetic diversity in *MLL-AF9* leukemia, we crossed the *Mll-AF9* knock-in mouse model^9^ with the five distinct inbred strains A/J, C57BL/6 (B6), 129S1/SvlmJ (129), NOD/ShiLtJ (NOD) and CAST/EiJ (CAST). We studied F1 hybrid mice heterozygous for *Mll*-*AF9* from these crosses versus the parental genetic background (Fig. 1A). The *MLL-AF9* parental strain has been maintained as it was historically, on a mixed B6 and 129 background. While the parental *Mll*-*AF9* strain has previously been demonstrated to develop leukemia around ∼6 months of age, detectable myeloid proliferation has been observed by 8 to 10 weeks of age^9,11^. Consistent with this observation, analysis of peripheral blood (PB) of parental *Mll-AF9* mice compared to *Mll*^WT^ littermates at 8 weeks of age showed significantly increased myeloid cell frequency with concomitant reduction in T cell frequency (Fig. 1B). This same phenotype was observed in *Mll-AF9* A/J F1, *Mll-AF9* B6 F1, *Mll-AF9* 129 F1 and *Mll-AF9* NOD F1 mice. In contrast, myeloid cell frequency in *Mll-AF9* CAST F1 mice was not significantly different from *Mll*^WT^ mice but instead a significant increase in B cell frequency and reduction in T cell frequency were observed. To determine whether this observation was based on baseline differences in the CAST genetic background, we examined PB composition in wild-type CAST mice versus the other strains used in this study. We found that wild-type CAST mice have no differences in PB composition compared to the other strains (Supp. Fig. 1), suggesting that this phenotype is a direct consequence of *Mll-AF9* expression.

**Figure 1.**
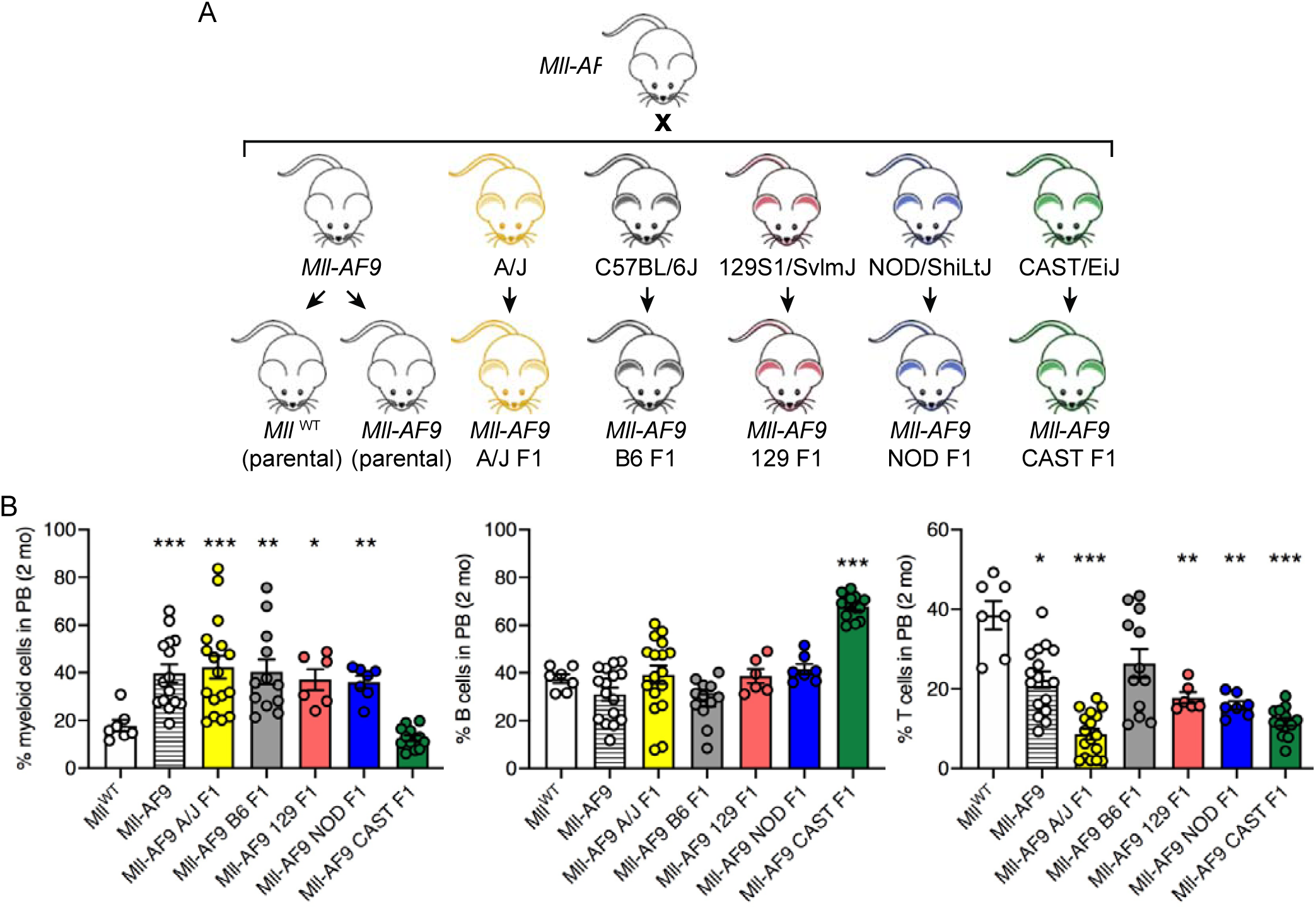
Genetic background-dependent differences in hematopoietic lineage composition in *Mll-AF9* knockin model. (A) Breeding strategy to create cohorts of genetically distinct F1 animals heterozygous for the well-characterized *Mll-AF9* knockin allele^9^. (B) Frequency of myeloid, B and T cells in the peripheral blood (PB) of indicated strains at 2 months of age. Dots represent individual mice (*Mll*^WT^, n=7; *Mll-AF9*, n=15; *Mll-AF9* A/J F1, n=17; *Mll-AF9* B6 F1, n=12; *Mll-AF9* 129 F1, n=6; *Mll-AF9* NOD F1, n=7; *Mll-AF9* CAST F1, n=13). Bars represent mean +/- SEM. \**P*<0.05, \*\**P*<0.01, \*\*\**P*<0.001 compared to *Mll*^WT^ parental values as determined by Brown-Forsythe ANOVA with Dunnett’s T3 multiple comparisons test.

Monitoring PB composition until pathology developed revealed that *Mll-AF9* CAST F1 mice maintained significantly reduced frequency of myeloid cells and increased frequency of B cells with aging compared to the parental *Mll-AF9* strain (Fig. 2A). In concordance with previous studies^11^, median survival in the parental *Mll-AF9* strain was 225 days. In contrast, *Mll-AF9* CAST F1 had a longer median survival (361 days; *P* = 0.079) and *Mll-AF9* A/J F1 and *Mll-AF9* 129 F1 that had significantly shorter median survival (172 days; *P* = 0.0071, and 178 days; *P* = 0.0179, respectively) (Fig. 2B). This data suggests that genetic background can alter the development and progression of leukemia caused by *Mll-AF9*.

**Figure 2.**
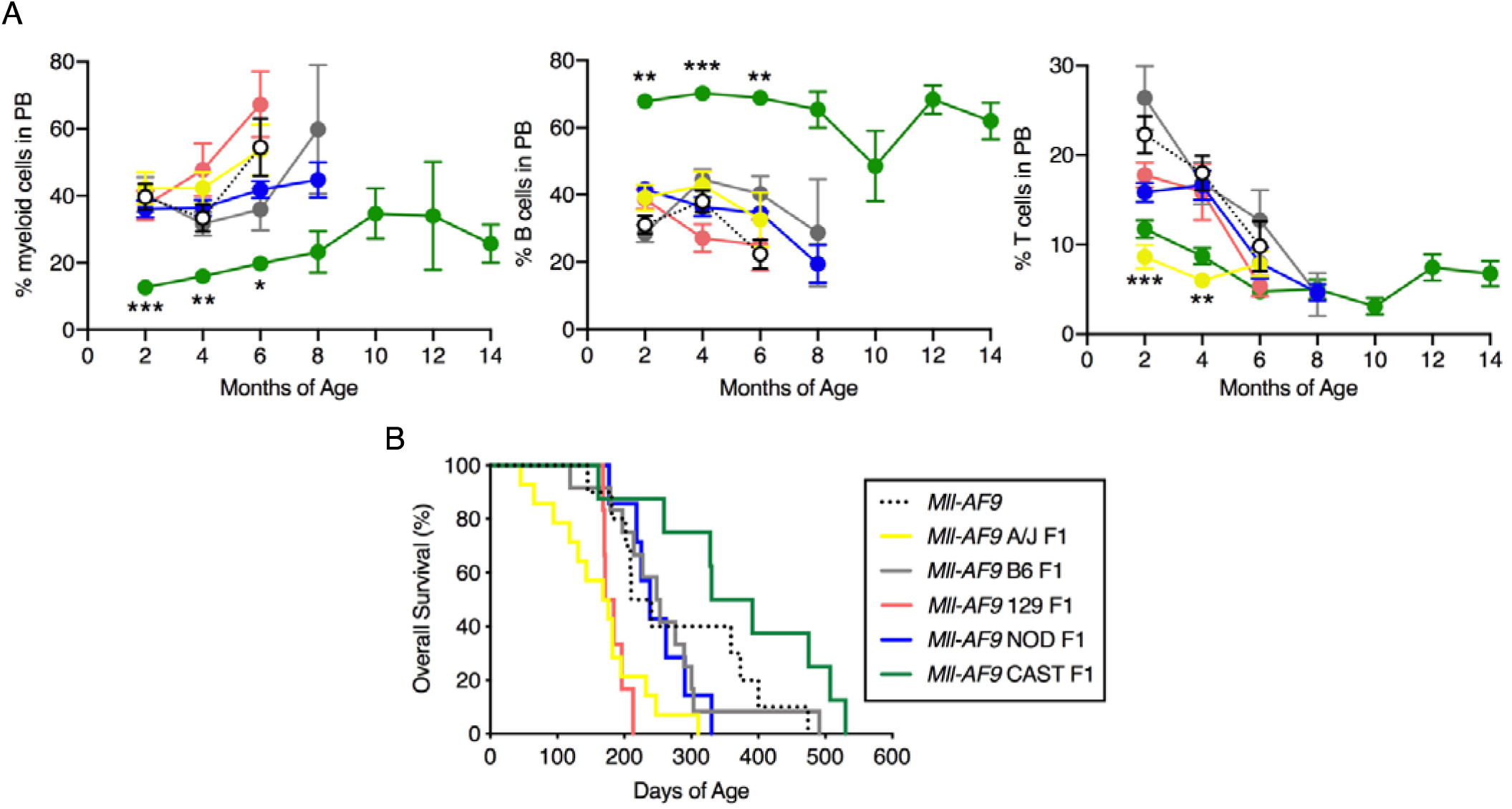
Genetic background-dependent progression to leukemia and overall survival mediated by *Mll-AF9*. (A) Frequency of myeloid, B and T cells in the PB of indicated strains from 2 months of age until moribund. Dots represent mean +/- SEM (*Mll-AF9*, n=15; *Mll-AF9* A/J F1, n=17; *Mll-AF9* B6 F1, n=12; *Mll-AF9* 129 F1, n=6; *Mll-AF9* NOD F1, n=7; *Mll-AF9* CAST F1, n=13). \*\**P*<0.01, \*\*\**P*<0.001 compared to parental *Mll-AF9* values as determined by Brown-Forsythe ANOVA with Dunnett’s T3 multiple comparisons test. (B) Overall survival of *Mll-AF9* mice (*Mll-AF9*, n=10; *Mll-AF9* A/J F1, n=14; *Mll-AF9* B6 F1, n=12; *Mll-AF9* 129 F1, n=6; *Mll-AF9* NOD F1, n=7; *Mll-AF9* CAST F1, n=8).

Characterization of the hematologic malignancies that developed in these strains also revealed genetic background-dependent distinctions. Consistent with previous studies^9,11^, parental *Mll-AF9* mice developed myelomonocytic AML with 100% penetrance (Fig. 3A) characterized by leukocytosis and thrombocytopenia (Fig. 3B), splenomegaly (Fig. 3C), >20% blasts in the BM (Fig. 3D), and abundant myeloid cell infiltration into the spleen and liver (Fig. 3D,E). Strains with shorter median survival (*Mll-AF9* A/J F1 and *Mll-AF9* 129 F1) developed either AML or an early and aggressive MPD-like disorder characterized by splenomegaly, <20% immature or blast-like cells in the BM, and high frequency of Gr1-expressing granulocytes in the BM (Fig. 3E). In the *Mll-AF9* CAST F1 strain exhibiting the longest median survival, 50% of mice were found to have ALL characterized by leukocytosis, splenomegaly, spleen and liver infiltration, and high frequency of B220^lo^ c-Kit^+^ blast cells in the bone marrow and spleen. While CAST mice have not been broadly studied in the context of leukemia or other cancer development, CAST F1 mice do have increased tumor growth in a model of neuroendocrine prostate carcinoma^15^, suggesting that the increased survival we have observed is not due to a general tumor-resistant genetic background. Of note, one individual *Mll-AF9* NOD F1 mouse was found in our study to develop MPAL characterized by leukocytosis, thrombocytopenia, spleen and liver infiltration, and bi-phenotypic B220^+^ CD11b^+^ blast cells in the bone marrow. As NOD mice are a polygenic model for autoimmune type 1 diabetes and exhibit aberrant immunophenotypes it is interesting to speculate how this influences the development of a bi-phenotypic leukemia.

**Supplemental Figure 1.**
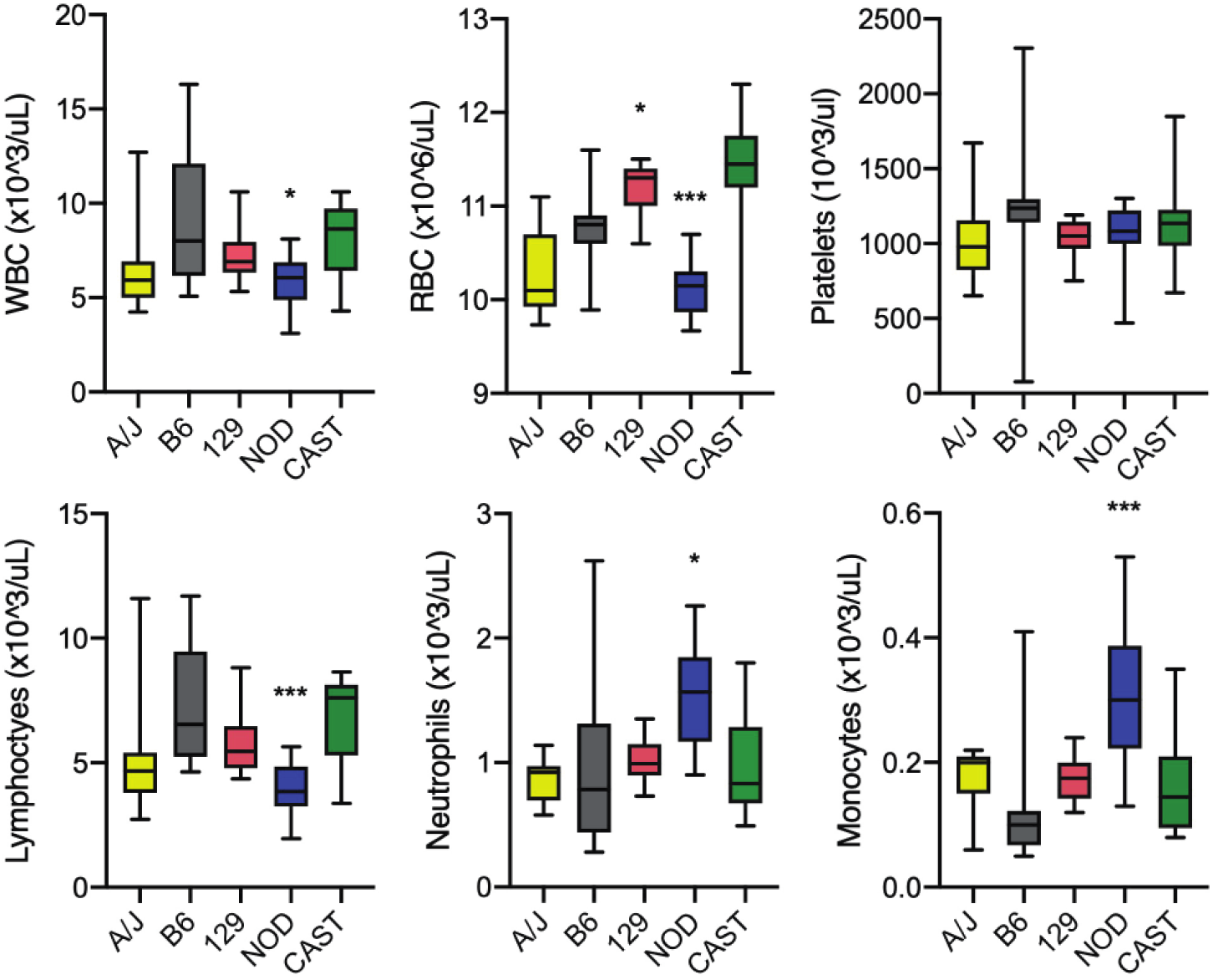
Complete blood count (CBC) values in inbred background strains used in this study. White blood cell (WBC), red blood cell (RBC), platelet, lymphocyte, neutrophil and monocyte counts in 6 month-old inbred A/J (n=13), B6 (n=15), 129 (n=20), NOD (n=16) and CAST (n=12) mice. Data obtained from Mouse Phenome Database^25^ (https://phenome.jax.org). Bars represent median, min to max. \**P*<0.05, \*\*\**P*<0.001 compared to inbred B6 values as determined by Brown-Forsythe ANOVA with Dunnett’s T3 multiple comparisons test.

**Figure 3.**
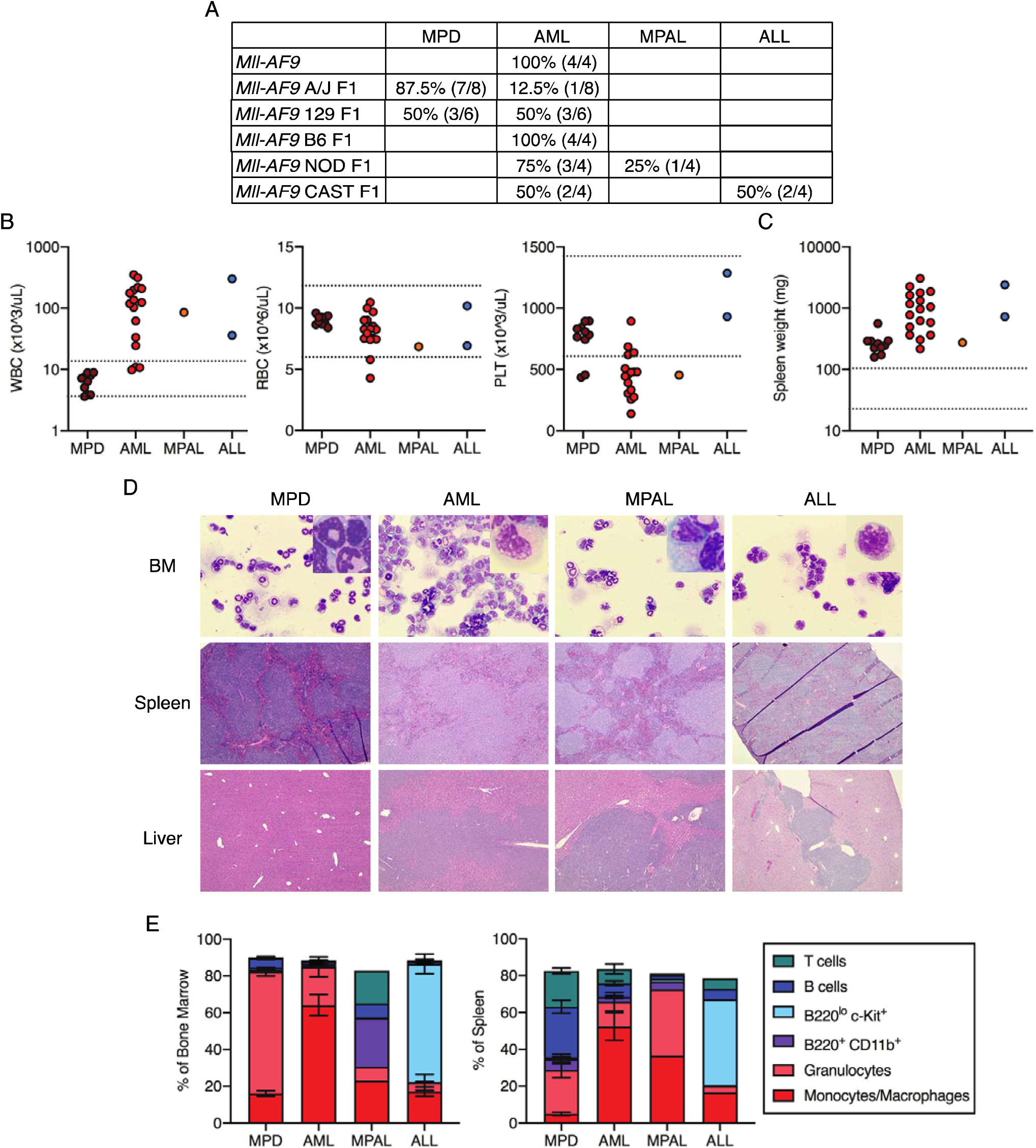
*Mll-AF9*-driven leukemia phenotype varies based on genetic background. (A) Frequency and numbers of mice diagnosed with myeloproliferative disorder (MPD), acute myeloid leukemia (AML), mixed phenotype acute leukemia (MPAL), and acute lymphoblastic leukemia (ALL). (B) WBC, RBC and platelet (PLT) counts in moribund mice. Dots represent individual mice. MPD, n=10; AML, n=16; MPAL, n=1; ALL, n=2. Dotted horizontal lines represent range of values considered to be normal in C57BL/6J mice^25^. (C) Spleen weight in moribund mice. Dots represent individual mice. MPD, n=10; AML, n=17; MPAL, n=1; ALL, n=2. Dotted horizontal lines represent range of values considered to be normal in C57BL/6J mice^25^. (D) Representative histological staining of bone marrow (BM) (Giemsa-stain, 40x, inset: 100x), spleen (H&E, 4x) and liver (H&E, 4x) in mice with MPD, AML, MPAL or ALL. (E) Flow cytometric analysis of indicated cell types in bone marrow and spleen of moribund mice. Bars represent mean +/− SEM of MPD, n=10; AML, n=14; MPAL, n=1; ALL, n=2.

By introducing genetic variation into the *MLL-AF9* knockin mouse model, our work has identified that disease latency and leukemia phenotype is significantly affected by heritable genetic variants segregating among common inbred strains, and suggests the presence of specific, dominant-acting modifier alleles in one or more strains. These findings support that genetic background differences may play a role in how and why leukemogenesis resulting from a common fusion oncogene can result in distinct etiology among different individuals. As epigenetic dysregulation is a critical driver of *MLL*-rearranged leukemia^16,17^, we posit that altered survival and leukemia phenotypes may be related to differences in epigenetic or chromatin state in genetically diverse mice. This is also supported by GWAS identification of single nucleotide polymorphisms in *ARID5B*, encoding part of the histone H3K9me2 demethylase complex, that modify risk for *MLL*-rearranged early childhood leukemia^4^. More broadly, apart from the presence of the fusion oncogene, data from our lab and others support that other factors strongly influence the specific outcome, including the cell type-of-origin^18-20^, the developmental stage and context in which the chromosome translocation occurs^21,22^, and the individual’s genetic background. Importantly, this study included five of the eight founder strains of the Collaborative Cross (CC) and Diversity Outbred (DO) mouse populations^23,24^, complementary resources that enable one to model human genetic diversity and map genetic modifiers that underlie phenotype differences in the population. Future studies will take advantage of these powerful tools to map the genetic determinants of leukemia susceptibility and phenotype, with the goal of identifying novel gene targets for the development of new therapies for *MLL*-rearranged leukemia.

## Methods

### Experimental animals

Kmt2a^tm2(MLLT3)Thr^/KsyJ (referred to as *Mll-AF9*, stock no: 009079) mice were obtained from, and aged within, The Jackson Laboratory. The *Mll-AF9* model was created on a 129P2/OlaHsd background, crossed with C57BL/6NCrl females for four generations, and has been maintained at The Jackson Laboratory since 2012 on a mixed C57BL/6 and 129S1/SvlmJ background by breeding with B6129PF1/J. The *Mll-AF9* original strain was crossed to the A/J, C57BL/6 (B6), 129S1/SvlmJ (129), NOD/ShiLtJ (NOD) and CAST/EiJ (CAST) strains to create F1 generation experimental mice. Male and female F1 progeny from each strain cross were included in the studies and monitored from 8 weeks of age until moribund. Female and male mice were analyzed for PB CBC data at 6 months of age. The Jackson Laboratory’s Institutional Animal Care and Use Committee (IACUC) approved all experiments.

### Peripheral blood analysis

PB was collected from mice via retro-orbital sinus and red blood cells were lysed before staining mature lineage markers: B220 (clone RA3-6B2), CD3e (clone 145-2C11), CD11b (clone M1/70), Gr-1 (clone RB6-8C5). Stained cells were analyzed on an LSRII (BD) and populations were analyzed using FlowJo V10. Differential blood cell counts were obtained from PB using an Advia 120 Hematology Analyzer (Siemens)

### Analysis of moribund mice

Moribund mice identified by declining health status were euthanized and PB, spleen, liver, and BM harvested. Single-cell suspensions of PB, spleen, and BM were analyzed by flow cytometry for mature lineage markers and c-Kit (clone 2B8), using an LSRII (BD) and populations were analyzed using FlowJo V10. Differential blood cell counts were obtained from PB using an Advia 120 Hematology Analyzer (Siemens). Cytospin preparations of whole BM MNCs were stained with May–Grunwald–Giemsa stain. Liver and spleens were fixed for 24 h in 10% buffered formalin phosphate, embedded in paraffin, and sections were stained with H&E. Histological images of stained BM, liver, and spleen were captured on a Nikon Eclipse C*i* upright microscope with SPOT imaging software (v.5.6).

### Statistical analysis

Overall survival, Log-rank (Mantel–Cox) test was performed on Kaplan–Meier survival curves. Statistical analysis of non-survival data was performed by Brown-Forsythe one-way ANOVA test followed by Dunnett’s multiple comparisons test. All statistical tests, including evaluation of normal distribution of data and examination of variance between groups being statistically compared, were assessed using Prism 8 software (GraphPad).

## Acknowledgements

This work was supported by National Institutes of Health (NIH), NCI Cancer Core Grant P30CA034196, and Eunice Kennedy Shriver National Institute of Child Health and Human Development (NICHD) T32HD007065 (K.Y.). This work was also supported by The V Foundation V Scholar award (J.J.T.) and grants from the Maine Cancer Foundation (J.J.T.). K.Y. is supported by an American Society of Hematology (ASH) Scholar Award and the Pyewacket Fund at The Jackson Laboratory. We thank Steve Munger, Jennifer SanMiguel, and members of the Trowbridge laboratory for helpful discussion and critical comments, and Rebecca Bell for experimental and laboratory support.

